# A network of face patches in human prefrontal cortex for social processing of faces

**DOI:** 10.1101/2025.05.05.652210

**Authors:** Asa Farahani, Mojan Izadkhah, Roza Hamidi, Elahe’ Yargholi, Gholam-Ali Hossein-Zadeh, Reza Rajimehr

**Author notes:** These authors contributed equally to this work (Asa Farahani, Mojan Izadkhah).

## Abstract

The human cerebral cortex contains localized regions for processing faces. These regions or patches, which are classically found in the occipito-temporal cortex, encode visual properties of faces. Using naturalistic movie-watching fMRI data from 176 human subjects and multivariate functional connectivity analysis, here we comprehensively characterize a novel network of four frontal face patches (FFPs) arranged dorsoventrally in the lateral prefrontal cortex. FFPs are strongly coupled with a face-selective region in the middle superior temporal sulcus, appear to be primarily involved in processing high-level social aspects of faces during movie-watching, and show partial correlations of activity with distinct cognitive networks. Activations in FFPs are correlated with the performance of subjects in a social cognition task. We further identify two groups of subjects who showed a remarkable difference in the topographical organization of FFPs. The discovery of FFPs provides new insights into the understanding of social processing in the brain.

## INTRODUCTION

Face recognition is a fundamental aspect of human vision, providing a wealth of vital information, including details about gender, race, age, identity, and emotional states [14]. The exploration of neural mechanisms underlying face processing has long been a central focus of cognitive neuroscience. This endeavor has led to the identification of a complex network of brain regions mediating this critical task [32, 37, 41, 42, 49, 68, 70, 103]. These regions are typically localized by blocked-design comparison of faces versus other categories. Based on this type of data, a seminal hierarchical model developed by Haxby and colleagues has played a significant role in characterizing the organization of the face-processing system. This model subdivided the face-processing system into two main components: the core and extended systems [41, 42]. The core system, including Fusiform Face Area (FFA) [73], Occipital Face Area (OFA) [68], and posterior Superior Temporal Sulcus (pSTS) [5], is involved in the visual analysis of faces. The extended system, comprising multiple areas outside the occipito-temporal cortex, complements the core system in extracting higher-level meanings from faces.

Extracting meanings from faces could involve prefrontal cortex, as prefrontal regions have been shown to play a significant role in high-level cognitive processing [62]. In fact, electrophysiological and neuroimaging studies in non-human primates have suggested the existence of face-processing regions in the prefrontal cortical regions. Scalaidhe et al. found a small population of highly face-selective cells in the prefrontal cortex of monkeys. These cells exhibited strong responses to faces while showing minimal or no responses to non-face stimuli [65, 66]. Rolls et al. also reported the existence of face-responsive neurons in the monkey’s orbitofrontal regions, which were involved in understanding facial expressions and social interactions [76, 77]. Using fMRI in macaque monkeys, Tsao et al. reported three regions in the prefrontal cortex, which demonstrated highly selective responses to face images [95]. The frontal face-responsive regions were also reported in marmosets while watching marmoset faces in movies [44]. Furthermore, Schaeffer et al. provided evidence for the role of marmoset’s frontal face regions in understanding social face stimuli [84]. These frontal regions in marmosets and macaques are also involved in the multimodal integration of species-specific facial and vocal information [25, 80].

Given the resemblance in brain organization across primates, it is reasonable to anticipate analogous functional zones within the human brain [44, 94, 106]. Supporting this notion, using fMRI, Tsao et al. demonstrated a face-selective patch near the right inferior frontal sulcus in three out of nine human participants, which was homologous to a region they had previously reported in macaques [94, 95]. Additionally, an fMRI study by Rajimehr et al. revealed a frontal face patch in the precentral gyrus near the frontal eye field (FEF) in both species [73]. Human studies have highlighted the involvement of prefrontal areas in the social understanding of faces. One study reported stronger activation of the lateral prefrontal regions in response to moving faces compared to static ones [64]. Moreover, applying Transcranial Magnetic Stimulation (TMS) over the inferior frontal gyrus resulted in a delay in associating a face with its adjective pairs, pointing to the role of the dorsomedial prefrontal cortex in the social impression formation [27].

Traces of prefrontal activation are seen in the group-average face maps, but the activation strength is comparably weaker in the prefrontal cortex compared to the occipitotemporal face-selective regions [106]. This relative weakness in prefrontal activations can be attributed to various factors. One possibility is that visually driven face-selective activations in the prefrontal cortex are genuinely weak because a dedicated neural machinery in the occipito-temporal cortex is already involved in the detailed analysis of facial features. Instead, the prefrontal regions are more involved in high-level processing of faces, such as decoding semantic meaning associated with faces, extracting social content of faces, and integrating facial information with other sensory cues. This idea is supported by studies showing that as opposed to FFA and OFA, which show visual field bias [55], frontal face patches do not exhibit such biases [64]. Methodological factors might also contribute to the weak face-related activations in prefrontal areas. For instance, the stimuli used in the fMRI localizer experiments usually include isolated and static face images which may not be robust enough to fully engage frontal regions [30]. Inter-subject variability in frontal areas introduces further complications, as spatial blurring caused by group-averaging tends to diminish statistical significance of activations. The blurring effect is exacerbated in frontal areas due to more variability in the location and quantity of sulci and gyri in these areas [4, 7, 15, 34]. Taken together, understanding the role of prefrontal cortex in face processing requires considering all these various factors and employing innovative methodologies.

Here we used naturalistic movie-watching fMRI data from the Human Connectome Project (HCP) from 176 healthy young subjects to explore the role of prefrontal areas in face processing. The movie-watching task can elicit diverse brain dynamics and allow us to capture more realistic brain activity patterns [39, 96], mitigating the problem of stimulus simplicity for driving the prefrontal activation. However, unlike well-controlled blocked-design stimuli, the movie-watching paradigm lacks structured categorically driven blocks, presenting an obstacle in identifying category-specific activations. We overcame this challenge using a data-driven, multivariate connectivity-based approach to localize the face-selective prefrontal regions [21]. Our study unveiled face-selective patches located within the right lateral prefrontal cortex, which were specifically connected to a face-selective area in the right middle Superior Temporal Sulcus (mSTS)—an important hub in the social processing network [19, 51]. We found that these areas were prominently involved in the social processing of faces, as their activations showed a strong correlation with the performance of subjects in the HCP social cognition task. We also investigated their functional properties and their intricate patterns of connectivity with other cortical regions. Furthermore, we took a novel approach for investigating the variability in the organization of prefrontal face patches across subjects and found two subtypes of individuals with distinct topological organization of these patches. These subtypes were also different from one another in both the underlying cortical structure and the activation patterns produced in the HCP social cognition task.

## RESULTS

In this study, we used a connectivity-based approach to identify face-selective regions in the prefrontal cortex. In this approach, we first defined the classic face-selective areas by evaluating the group-average activation map (averaged beta map) for the face category, obtained by statistically comparing the blocks of faces versus the blocks of bodies, places, and tools in the HCP *N*-back working memory task (Figure 1A,B). This map revealed well-established hotspots of face activations in the occipito-temporal cortex. Subsequently, we defined eight cortical face-selective areas using a thresholding method applied to the face activation map, considering the top 1% of vertices/voxels with the highest face-selectivity. These face-selective areas consisted of three areas in the left hemisphere, including FFA, pSTS, and Medial Face Area (MFA), and five areas in the right hemisphere, including FFA, OFA, pSTS, mSTS, and MFA (Figure 1C,D). The discrepancy in the number of identified areas between the two hemispheres corroborates previous studies reporting the superiority of the right hemisphere involvement in face processing [24, 26, 48, 56, 81, 88]. Additionally, an examination of unthresholded group-average face activation map revealed some weak activations in the prefrontal cortex, more clearly seen in the right hemisphere (highlighted by the white rectangle in Figure 1B). This prefrontal activation, while present, appeared notably less pronounced compared to the robust face-related activity in the occipito-temporal cortex.

**Figure 1.**
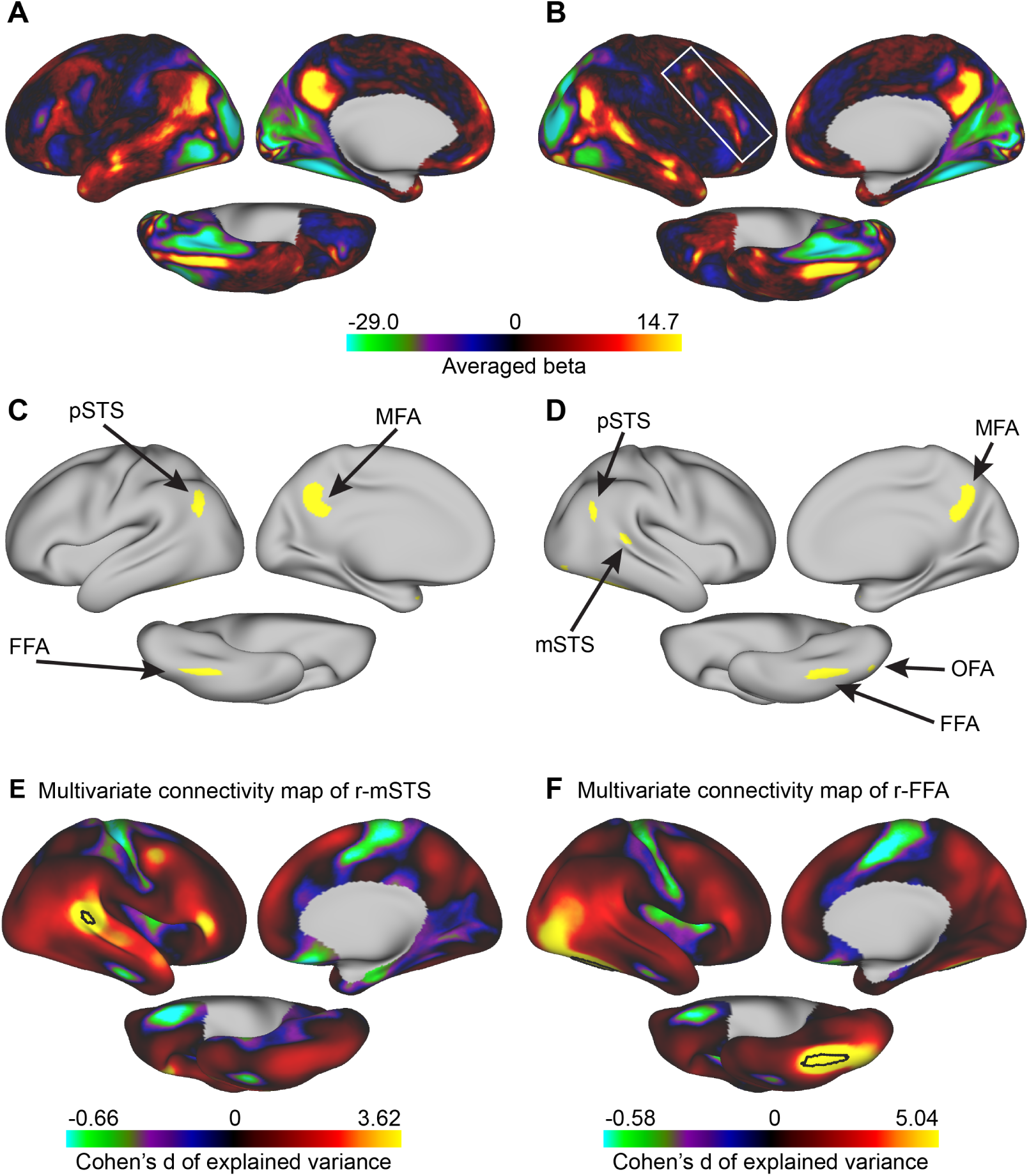
Classic face-selective areas and their multivariate cortical functional connectivity patterns. Brain cortical maps are shown on lateral, medial, and ventral views of inflated fs-LR surface. (A) Left hemisphere’s group-average activation map from the HCP-S1200 package for the contrast of faces vs. other categories (bodies, places, and tools) in the HCP *N*-back working memory task. (B) Similar to (A), but for the right hemisphere. The white box highlights the prefrontal regions activated during the face localizer task. (C) Classic face-selective areas in the occipitotemporal cortex. These areas include FFA, pSTS, and MFA in the left hemisphere (D) and FFA, OFA, pSTS, mSTS, and MFA in the right hemisphere. These areas are identified by thresholding the group-average beta coefficient map to include 1% of the vertices/voxels in the CIFTI grayordinate space (913 out of 91, 282 vertices/voxels) with the largest beta values for the faces vs. other categories contrast. (E) Multivariate connectivity map of the right mSTS (black outline) with ipsilateral cortical regions. (F) Multivariate connectivity map of the right FFA (black outline) with ipsilateral cortical regions. Unlike mSTS, FFA does not show strong connectivity with prefrontal regions.

In the next step, we obtained a multivariate connectivity map for each face-selective area in the occipito-temporal cortex. To derive these maps, we used the HCP 7T movie-watching fMRI data from 176 healthy young adults (https://www.humanconnectome.org/study/hcp-young-adult). Multivariate connectivity was assessed by predicting the mean time-course of activity in each occipito-temporal face area (target area) based on the pattern of activity in other cortical regions (seed regions) in the ipsilateral cortex (Supplementary Figure S1). To accomplish this, we employed the whole-cortex surface-based searchlight analysis [50]. The multivariate connectivity approach has higher sensitivity for uncovering the functional interactions between areas compared to the univariate correlation-based methods [105].

A regression analysis was used to predict the mean time-course of activity in a target area based on the pattern of activity in all searchlight regions of interest (ROIs) (Supplementary Figure S1). Instead of ordinary regression, we employed Partial Least Squares Regression (PLSR) which minimizes the collinearity problem often seen in regression analyses [2, 3, 60]. Details of PLS model’s training and testing procedures are provided in *Methods* and briefly described here. For each searchlight ROI, a PLS model was constructed/trained using a subset of movie data to find a mapping between the mean time-course of activity in the target area and the time-courses of activity in 100 vertices within the ROI. For this mapping, PLSR searches for a set of components that perform a simultaneous decomposition of predictors and dependent variables. Next, the trained model was used to predict the mean activity of the target area in the test movie data. The predicted activity was then compared with the actual activity, and an explained variance value was estimated. A higher explained variance value indicates that the searchlight ROI has better prediction power and possibly stronger multivariate connectivity with the target area. This value was assigned to the central vertex of the searchlight ROI. Performing this procedure for all searchlight ROIs resulted in a cortical map of explained variances (a multivariate connectivity map). The multivariate connectivity maps were obtained for each classic face-selective area in each individual subject.

Our data-driven approach revealed notable multivariate connectivity between patch-like areas of the lateral prefrontal cortex and the mSTS face area (Figure 1E). The group-average multivariate connectivity of prefrontal areas with other face-selective areas was comparatively weaker (Supplementary Figure S2). This different pattern of multivariate connectivity was exemplified by the right FFA’s group-average connectivity map, which did not exhibit distinct hotspot-like regions in the lateral prefrontal cortex (Figure 1F). Given the established role of mSTS in social face perception [19, 51], its connectivity with the prefrontal patches implies that these patches may be involved in social cognitive processes. To explore the face-processing network within the prefrontal cortex, we focused on the group-level multivariate connectivity map of mSTS. Using a liberal threshold for the map, we defined a mask that encapsulated prefrontal patches demonstrating strong connectivity with mSTS (Figure 2A). Examining this delineated prefrontal territory in individual subjects revealed the presence of three prefrontal face patches in the majority of subjects, albeit with some variation in the topology of patches across subjects (see Supplementary Figure S3 for all subjects’ data and Figure 2B for a subset of randomly selected subjects). Notably, the most dorsal and the most ventral face patches displayed consistent positioning across subjects, while the positioning of the central patch varied considerably.

**Figure 2.**
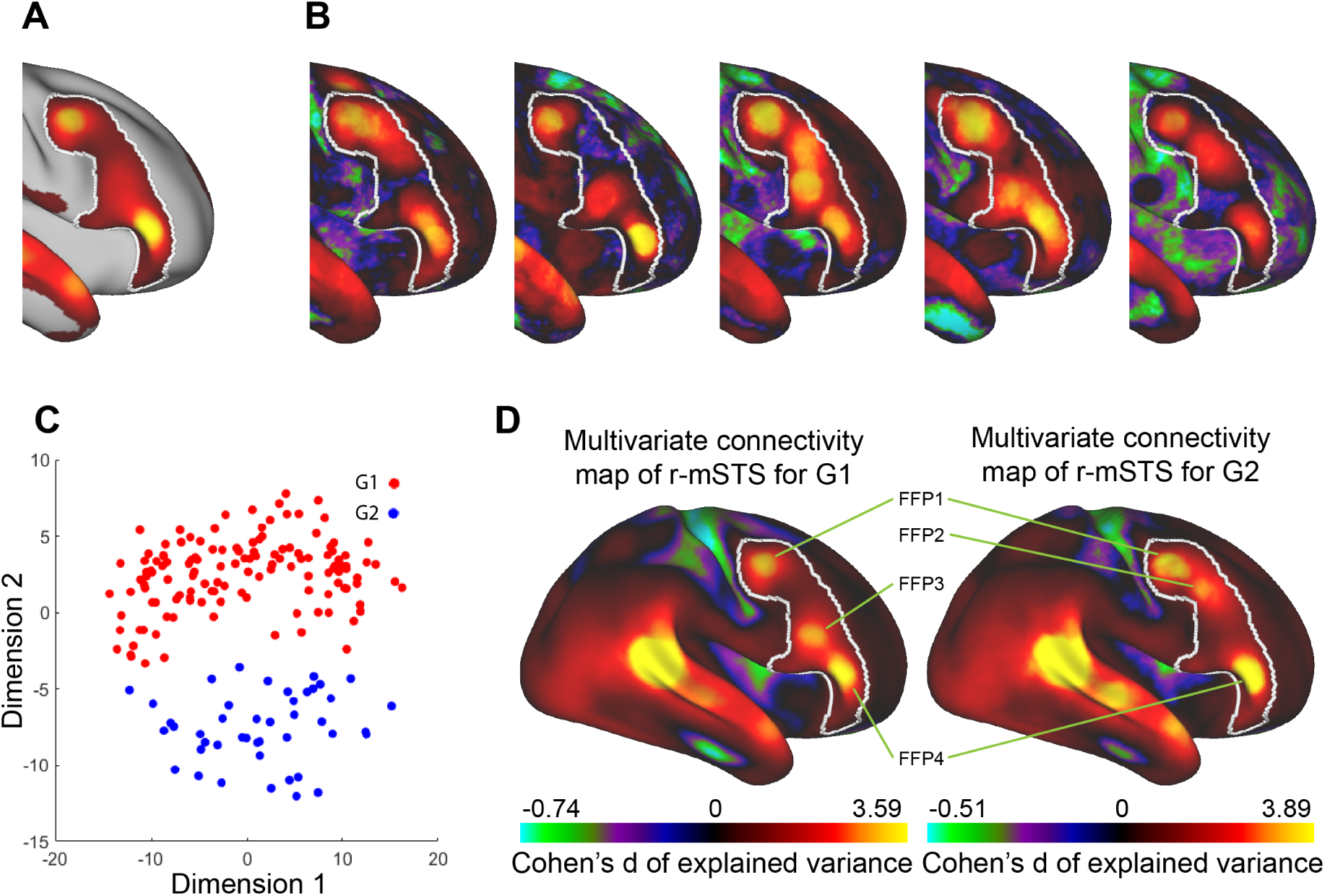
Group differences in the organization of Frontal Face Patches (FFPs). All brain cortical maps in this figure are shown on the lateral view of the right hemisphere’s inflated fs-LR surface. (A) A prefrontal mask (white outline) containing patches of high connectivity with the right mSTS. (B) Individual connectivity maps of the right mSTS with regions in the predefined mask for five randomly selected subjects. (C) A low-dimensional representation of the connectivity patterns within the mask across all 176 subjects (each subject is presented as a dot). Based on a clustering algorithm, two groups of subjects (G1 and G2) are identified. Red dots indicate G1 subjects (*N* = 132), while blue dots represent G2 subjects (*N* = 44). (D) The group-average connectivity map of the right mSTS shows distinct profiles for G1 (left) and G2 (right). Both groups exhibit FFP1 and FFP4 in similar locations. However, FFP2 is unique to G2, and FFP3 is unique to G1. The prefrontal mask’s boundary is shown in white.

Variations in the topological distribution of face-selective patches within the prefrontal cortex raised a crucial question: Does the variability in the location of the central patch across subjects happen randomly, or does it have a specific pattern that could hint at distinct subtypes of subjects? To address this question, we employed dimensionality reduction via Isomap [93], followed by DBSCAN (Density-Based Spatial Clustering of Applications with Noise) clustering algorithm to classify subjects based on the organization of face patches in the prefrontal cortex. This analysis revealed the presence of two distinct subtypes within our study cohort: Group 1 (G1), consisting of 132 subjects (81 females, 51 males), and Group 2 (G2), consisting of 44 subjects (25 females, 19 males) (Figure 2C). By examining each subtype’s averaged multivariate connectivity map of mSTS and applying a thresholding procedure, we identified four Frontal Face Patches (FFPs), denoted as FFP1, 57FFP2, FFP3, and FFP4. The most dorsal and the most ventral patches, FFP1 and FFP4, were consistently present in all subjects. FFP2 was present only in G2, whereas FFP3 was present only in G1 (Figure 2D). Anatomically, FFPs were arranged dorsoventrally in the lateral prefrontal cortex. Based on Glasser’s parcellation [34], FFP1 intersected with areas 55b and FEF. FFP2 was located within area 8C. FFP3 was at the junction of areas IFSa, IFSp, and 44. FFP4 primarily overlapped with area 45, while also extending into FPO5 and IFSp.

To explore potential structural or functional correlates for these differences among the identified subtypes, we first compared the curvature (gyrus-sulcus) maps derived from structural images of the two subject groups. Figures 3A and 3B show the average curvature maps for G1 and G2, respectively. By statistically comparing the maps using Threshold-Free Cluster Enhancement (TFCE) [91], we observed marked differences in cortical curvature adjacent to FFPs (Figure 3C). In the vicinity of FFP2, cortical curvature was less negative in G1 than in G2, indicating that the underlying tertiary sulcus was deeper in G2. Conversely, in the vicinity of FFP3, cortical curvature was more positive in G2 than in G1, indicating that the underlying tertiary gyrus was more protruded in G2 (Figure 3C). These findings highlights a fundamental structural distinction between the two subject groups at the location of FFPs. Moreover, group-average fMRI maps of the HCP social cognition task revealed a robust social activation at the location of FFP1, FFP3, and FFP4 in G1 and a robust social activation at the location of FFP1, FFP2, and FFP4 in G2 (Figure 3D,E). The TFCE analysis showed that differences between the maps of the two subject groups were largely confined to the location of FFP2 and FFP3 (Figure 3F). These findings had two important implications: (1) they confirmed differences in the functional organization of prefrontal cortex between the two subject groups using an independent dataset, and (2) they provided further evidence for a direct involvement of FFPs in the processing of social interactions.

**Figure 3.**
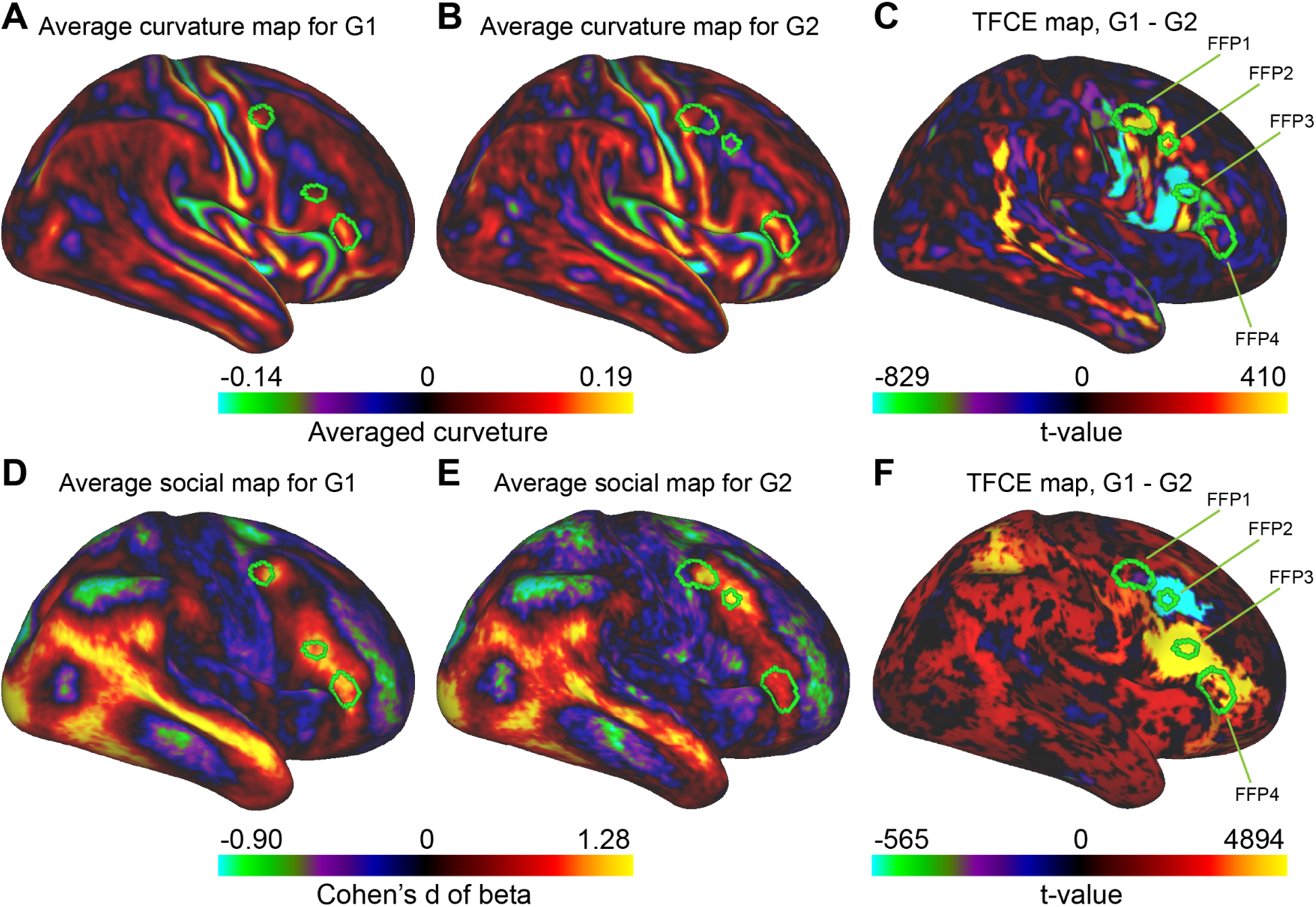
Structural and functional distinctions of FFPs between the two subject groups. All cortical maps in this figure are shown on the lateral view of the right hemisphere’s inflated fs-LR surface. (A) Group-average curvature map for G1. (B) Group-average curvature map for G2. (C) A distinct curvature pattern adjacent to FFPs is observed in the comparison between the two subject groups (t-value map from TFCE analysis). Notably, in G2, the underlying tertiary sulcus near FFP2 is deeper, and the tertiary gyrus near FFP3 is more protruded. (D) Group-average Cohen’s d effect size map of social cognition task for G1. A robust activation is observed in the vicinity of FFP1, FFP3, and FFP4. (E) Group-average Cohen’s d effect size map of social cognition task for G2. A robust activation is observed in the vicinity of FFP1, FFP2, and FFP4. (F) The difference in social cognition activation maps across the two groups (t-value map from TFCE analysis) emphasizes functional variations corresponding to distinct FFP configurations. In all panels, regions outlined with green borders highlight FFPs.

In the next step, we characterized the basic fMRI response properties of FFPs. Firstly, we calculated the mean activity of each FFP across subjects during the passive movie-watching (Figure 4A). FFP1 and FFP4 were present in both subject groups. Thus, the mean time-course in these two patches was calculated across all subjects after merging FFP masks across G1 and G2. However, for FFP2 and FFP3, we opted to calculate group-specific mean time-course within G2 and G1, respectively, as each patch was only available in a single subtype of subjects. The Pearson correlation analysis revealed a strong similarity between the mean time-courses of all FFPs (Figure 4B). We then examined the response variability within each patch by assessing the standard deviation of its mean time-course. This analysis exhibited the highest response variability in FFP2 (Figure 4C).

**Figure 4.**
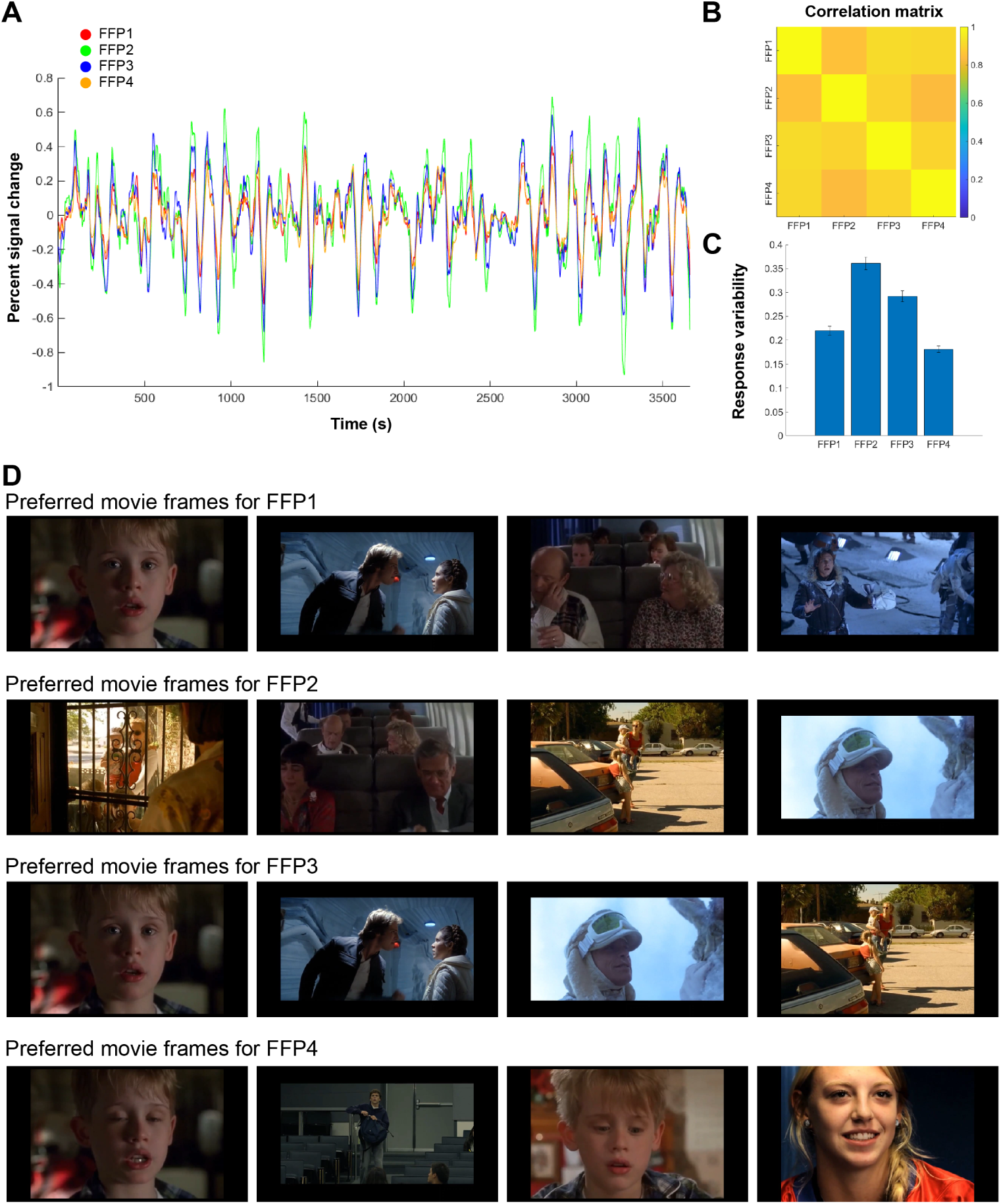
Response properties of FFPs during the movie-watching task. (A) Mean time-course of activity for each FFP, averaged across subjects having the corresponding patch. The functional data is composed of four movie-watching scans, totaling 3, 655 TRs. (B) The heatmap represents the interconnectivity of FFPs, measured via Pearson correlation of their mean time-courses. (C) The bars show the averaged standard deviation of FFP’s time-courses. Error bars indicate one standard error of mean across vertices. (D) For each FFP, the first four preferred movie frames are shown. The preferred movie frames of an FFP are the ones that produce the highest response in that patch. To find the preferred movie frames of a patch, we first obtain the mean time-course of activity across vertices of the patch, then the peaks of response are detected using the peak-detection algorithm of Matlab. The resulting time-points are sorted based on the magnitude of response, and 50 time-points with the highest response are selected. These time-points are then reordered based on an averaged activity in a 5-second window around each time-point. The movie frames corresponding to these time-points are obtained after considering a standard hemodynamic lag of 4 seconds for BOLD response [61]. In the figure, the preferred movie frames are ordered from left to right.

Considering the mean time-course activity of each patch, we unveiled the preferred movie frames associated with each FFP using a ‘reverse-correlation’ approach. In this approach, the movie frames that produced the highest response in each patch were identified. These movie frames could help formulate hypotheses about the function of these patches. The top four preferred movie frames in all FFPs included faces within a naturalistic context (Figure 4D). This result was in line with the result of multivariate connectivity analysis, supporting the role of FFPs in face processing.

We further explored the functional role of FFPs by assessing semantic category labels provided by the HCP dataset for every second of the movie clips. These annotations, generated using the WordNet semantic taxonomy [45], provided a set of descriptive terms (600 distinct object and action categories) summarizing the semantic content of the movie frames. For every label, we constructed a regressor/variable for the entire movie session. The regressors were binary vectors indicating the presence or absence of labels. We then correlated the regressors with the mean time-course of activity in each FFP after considering a standard hemodynamic lag of 4 seconds for BOLD response [61]. As shown in Figure 5, the terms “man” and “woman” had the highest correlation with the activity in FFPs. The term “talk” also showed a high correlation in all FFPs except in FFP2. These findings highlighted that the responses in FFPs were indeed highly selective to faces. In addition, the high response to conversations implied the possible role of FFPs in integrating faces with social cues.

**Figure 5.**
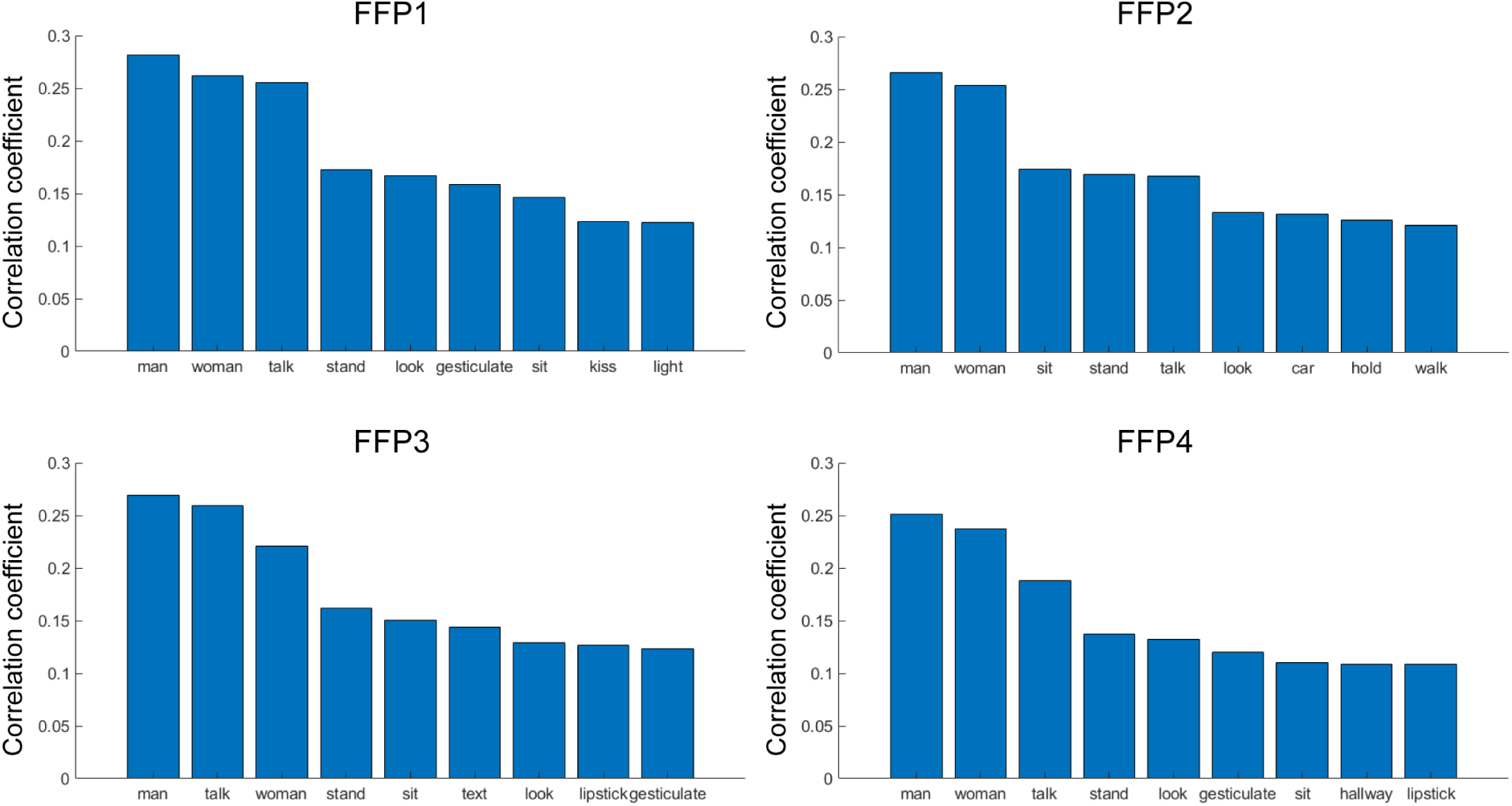
FFPs respond to facial and social information during the movie-watching task. The mean time-course of activity in each FFP is correlated with 600 word regressors after considering a standard hemodynamic lag of 4 seconds for BOLD response [61]. The regressors are defined by the semantic labels assigned to each second of the movie clips. The top nine semantic terms with the highest correlations are shown.other patches (Figure 6A-D) [59]. The parcel-wise partial correlation maps were also obtained by considering the mean time-course of activity in each parcel of Glasser’s parcellation. This analysis resulted in four values per parcel, each corresponding to the partial correlation of that parcel with an FFP. These four values underwent a simple comparative analysis – a winner-take-all procedure, which determined the FFP with preferential connectivity to the parcel. The preferential connectivity of all parcels with FFPs was then visualized as a map on inflated cortex (Figure 6E) and flat patch (Figure 6F) of the right hemisphere. To highlight parcels with strong connectivity, this map showed 90 out of 180 parcels with the highest partial correlation values.

As a complementary analysis, we looked into the univariate functional correlation between the mean time-course of activity in FFPs and the time-courses of all cortical vertices in the right hemisphere. The vertex-wise correlation map is shown in Supplementary Figure 4. As expected, we observed strong positive correlations in occipito-temporal cortical areas, precisely where classic face-selective regions resided. In particular, the Temporo-Parietal Junction (TPJ) showed the highest positive correlation value. TPJ is renowned for its involvement in social processing [31, 83, 85], reinforcing the crucial role of FFPs in social face perception. Regions within the superior temporal sulcus and the medial pre-frontal cortex, which play a pivotal role in the social brain network [11, 36, 82, 86], also demonstrated prominent positive correlations with FFPs.

All the analyses thus far strongly supported the idea that FFPs are involved in the processing of facial and social information. However, it is still unclear whether these patches contribute equally to the social face processing or each patch plays a specific role in this process. To address this question, we used a partial correlation analysis to determine the cortical areas that were preferentially connected to each patch. Examining these areas could provide insights into the specific function of each patch. To obtain the partial correlation maps, the mean time-course of activity in each patch was correlated with the time-courses of all ipsilateral cortical vertices after removing the correlations induced by

Through the partial correlation analysis, we found that each patch was preferentially connected to specific cortical parcels. FFP1, located in areas FEF and 55b, showed preferential connectivity with parcels MST, TPOJ2, PSL, PFcm, SCEF, 6ma, p32pr, FOP1, and FOP4. Some of these parcels, including FEF and SCEF, are involved in the control of eye movements [6, 67]. Another set of parcels, including 55b and PSL, are regions in the right hemisphere, which are involved in the processing of social interactions [71]. Thus, FFP1 may extract dynamic social cues to guide eye movements. FFP2, located in area 8C, showed preferential connectivity with parcels V3, V3A, V3B, V3CD, V4, V4t, V6A, V8, PIT, LO1, LO2, LO3, MT, TPOJ1, TPOJ3, PGp, IP0, FFC, TE2p, VVC, VMV1, VMV2, VMV3, PHA1, PHA2, PHA3, ProS, DVT, 7m, 7Am, 7Pm, 8Ad, PCV, POS1, v23ab, LIPd, STV, STSva, STSda, PEF, JFJp, and IFSp. The majority of these parcels were located within visual cortex and scene-selective regions. Thus, FFP2 may contribute to the integration of facial information with the background scene context. The selective connectivity of FFP2 with visual regions can explain the observed greater variability in its response to movies compared to other FFPs (Figure 4C). The source of this variability could be transient changes of visual stimuli in movies. FFP3, located in IFJa, showed preferential connectivity with parcels A1, A4, A5, STGa, TA2, 52, 47m, MBelt, RI, RBelt, LBelt, and OP4. These parcels were predominantly located within auditory cortex. Thus, FFP3 may integrate facial and acoustic information. Previous studies in monkeys have also reported regions in the prefrontal cortex for integrating facial and vocal information [46, 78, 79, 92]. FFP4, located in area 45, showed preferential connectivity with parcels 31pv, 31pd, PFM, PGi, STSdp, STSvp, TGd, TGv, 23d, 10pp, 44, 47s, 47I, FOP5, 8Av, 8BL, 9a, 9m, 9-46d, 9p, SFL, and s6-8. These parcels were mainly located within the default mode network. Given the role of default mode network in semantic processing [52], FFP4 may be particularly involved in social semantic processing of faces.

**Figure 6.**
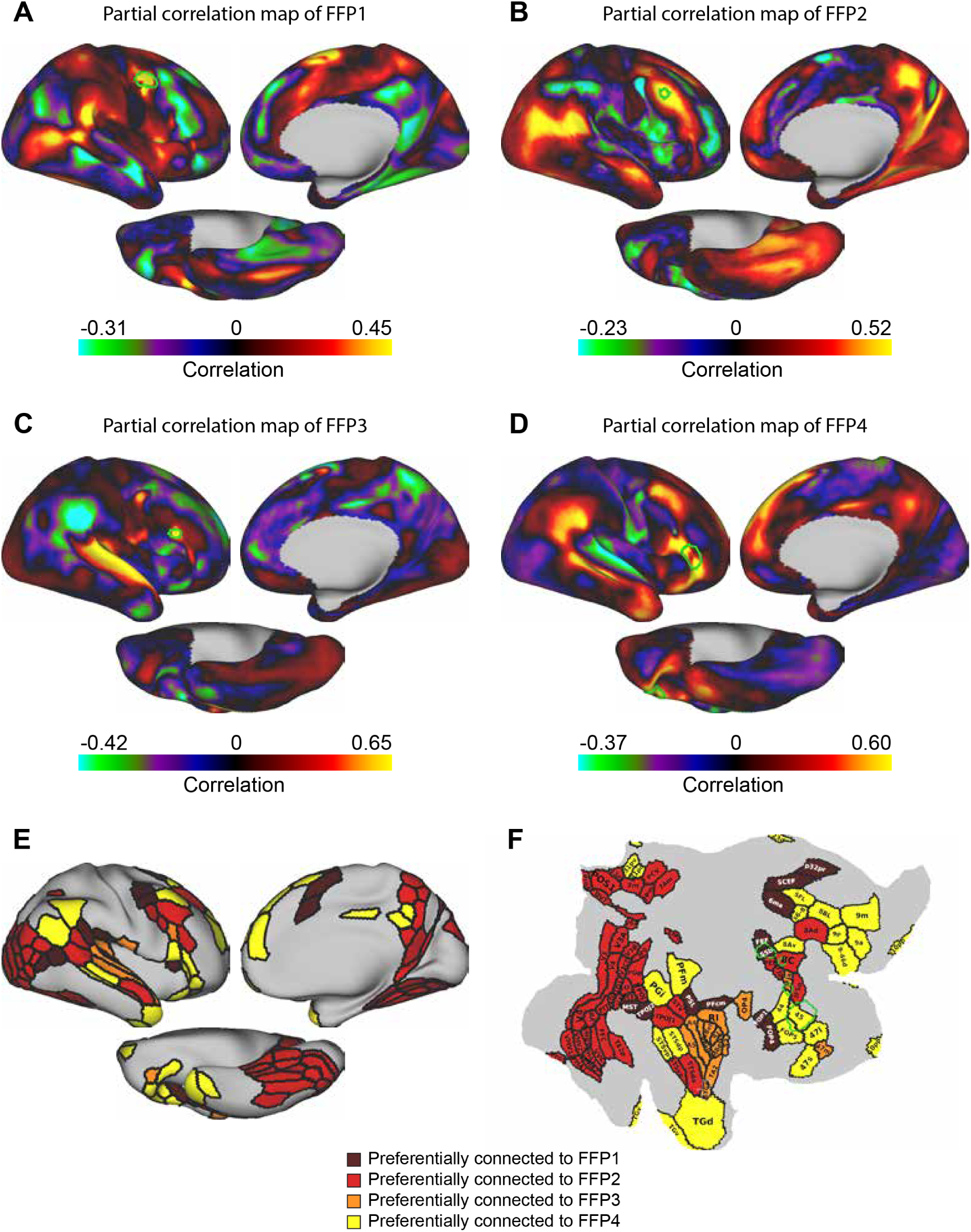
Unique cortical functional connectivity patterns of FFPs. (A-D) The partial correlation coefficient maps for FFPs are shown on lateral, medial, and ventral views of the right hemisphere’s inflated fs-LR surface. (E,F) Using a winner-take-all procedure, the cortical parcels preferentially connected to each FFP are obtained. The parcels are shown on the inflated (E) and flattened (F) fs-LR surface, and are labeled based on the multi-modal parcellation of the cerebral cortex [34]. The green outlines indicate FFPs.

To further justify these results, we qualitatively compared the associated parcels of FFP2, FFP3, and FFP4 with the HCP group-average maps from the task fMRI data. The associated parcels of FFP2 overlapped with scene-selective regions identified based on the contrast of scenes versus all other categories (faces, bodies, and tools) in the HCP *N*-back working memory task (Figure 7A,B). The associated parcels of FFP3 overlapped with the auditory cortex identified based on the contrast of auditorily presented stories versus baseline in the HCP language task (Figure 7C,D). Lastly, the associated parcels of FFP4 closely matched with regions involved in the semantic processing. These regions were identified based on the contrast of story versus math in the HCP language task (Figure 7E,F).

**Figure 7.**
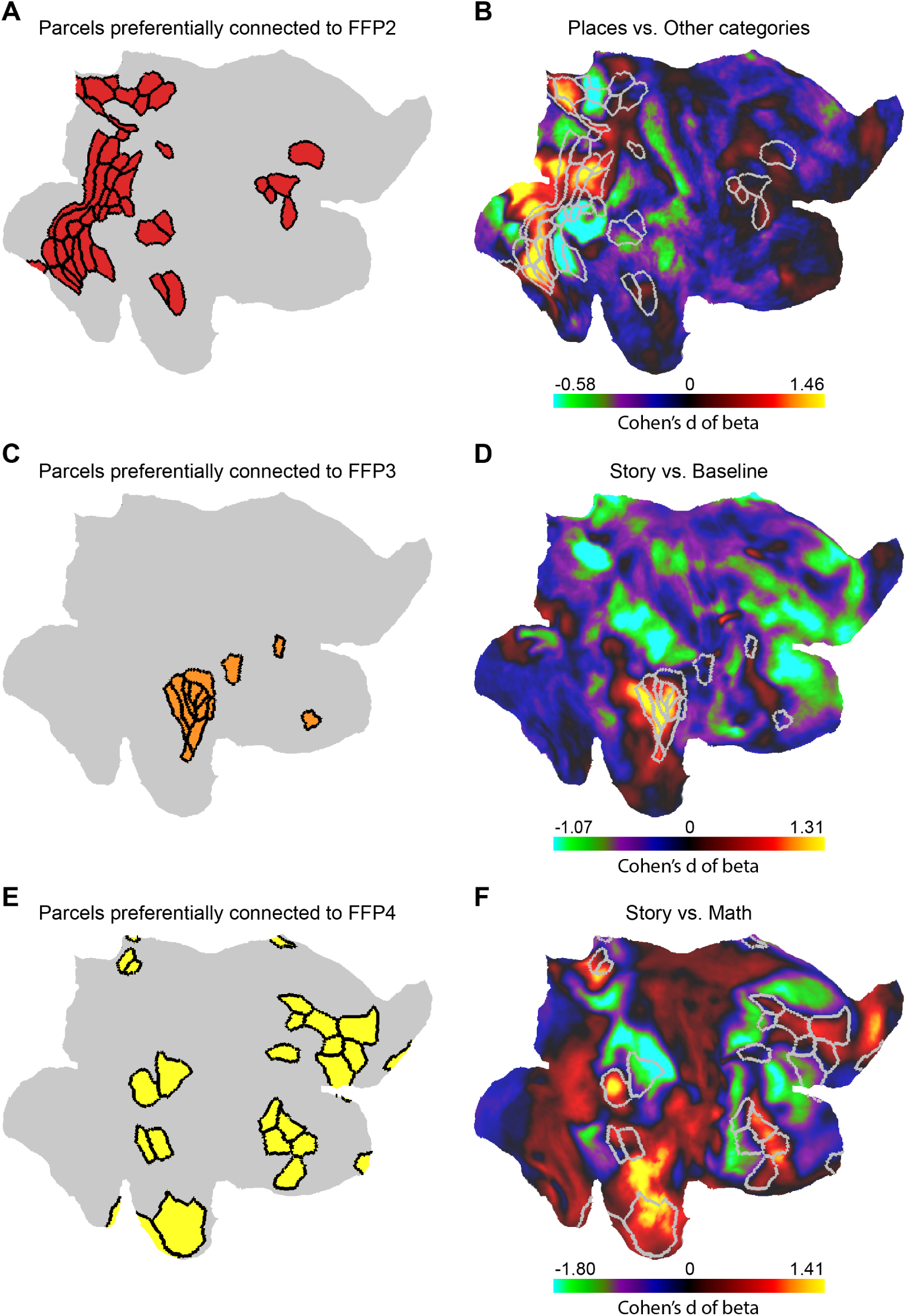
Topographic correspondence between preferentially connected parcels of FFPs and task fMRI activation maps. Brain cortical maps in all panels are shown on the right hemisphere’s inflated fs-LR surface. (A) Parcels preferentially connected to FFP2. (B) Group-averaged Cohen’s d effect size map for the contrast of places vs. other categories (faces, bodies, and tools) in the HCP *N*-back working memory task. (C) Parcels preferentially connected to FFP3. (D) Group-averaged Cohen’s d effect size map for the contrast of story vs. baseline in the HCP language processing task. (E) Parcels preferentially connected to FFP4. (F) Group-averaged Cohen’s d effect size map for the contrast of story vs. math in the HCP language processing task. In panels (B), (D) and (F), gray borders illustrate the parcels with high connectivity preference to each FFP.

To underscore the role of FFPs in social processing, we examined their activation in relation to behavioral performance in a social task. In this regard, we used data from the HCP social cognition task, which included social activation maps and social behavioral measures from 1, 048 subjects. In this task, subjects viewed short video clips containing animated shapes that could display random movements or movements implying social interactions, resembling scenarios that might evoke a “Theory of Mind” (TOM) interpretation. Subjects were instructed to report the content of clips as either TOM, Random, or Unsure. Figure 8A shows all potential combinations of stimuli and subjects’ responses. Figure 8B shows the quantitative distribution of subjects’ responses regardless of the stimuli presented. It was evident that Unsure responses were less prevalent. TOM and Random responses peaked near 50%, with a small bias for reporting TOM.

**Figure 8.**
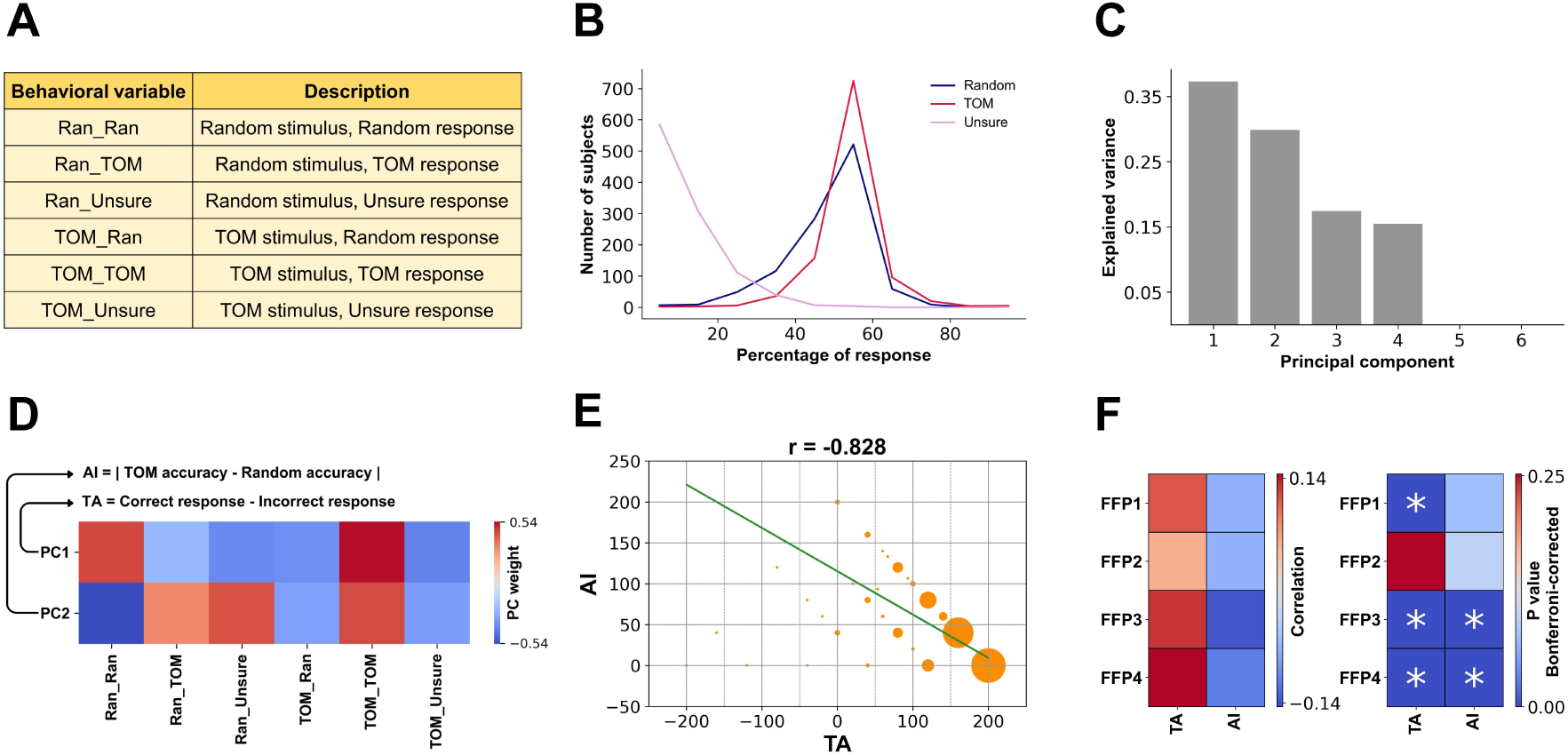
Relationship between the activations in FFPs and behavioral performance in the HCP social cognition task. (A) All potential stimulus-response combinations during the HCP social cognition task. For any stimulus type (Ran or TOM), subjects’ responses can be Ran, TOM, or Unsure. (B) Quantitative distribution of all 1, 048 subjects’ responses regardless of the stimulus type. (C) PCA is applied to the six behavioral measures. The bars show the explained variance value per component. The first two components account for up to 67.11% of the total variance. (D) Weights of these two components on the original six variables. Based on the profile of PCA weights, we characterize two measures of subjects’ performance: Total Accuracy (TA) defined as the difference between correct and incorrect responses, and Accuracy Imbalance (AI) defined as the absolute difference between accuracies for TOM and random conditions. (E) Inverse relationship between TA and AI. For each data-point in the scatter plot, the size of circle indicates the number of subjects. The green line is the linear regression line. (F) TA positively correlates with activations in FFPs during the social cognition task (significant for FFP1, FFP3, and FFP4; Bonferroni-corrected *p <* 0.05), while AI shows a negative correlation (significant for FFP3 and FFP4; Bonferroni-corrected *p <* 0.05).

We used Principal Component Analysis (PCA) on the six behavioral measures of all subjects to uncover latent variables explaining most of the variance in responses. This analysis revealed that the first two principal components accounted for up to 67.11% of the variance (Figure 8C). The weights of these components on the original six variables are shown in Figure 8D. The analysis of weights led us to define two new variables related to the first two principal components. The first variable, named Total Accuracy (TA), was the difference between correct and incorrect responses. The second variable, named Accuracy Imbalance (AI), was the absolute difference between accuracies for TOM and Random conditions. Note that there was an equal number of TOM and Random blocks during scans, and therefore, a higher AI indicated a worse performance. As shown in Figure 8E, AI was indeed inversely related to TA (Spearman correlation coefficient = *−*0.828). Thus, TA and AI were both measures of subjects’ performance. With respect to the subject groups, the mean TA was slightly higher in G1 than G2 (159.3 in G1 and 150.3 in G2), though the difference was not statistically significant (*p >* 0.05, t-test).

TA was positively correlated with activations in FFPs during the social cognition task (Figure 8F). The positive correlation was significant for FFP1, FFP3, and FFP4 (Bonferroni-corrected *p <* 0.05). Conversely, AI had a negative correlation with activations in FFPs (Figure 8F). The negative correlation was significant for FFP3 and FFP4 (Bonferroni-corrected *p <* 0.05). These results indicated that enhanced activation within these patches could lead to improved performance in a social task.

## DISCUSSION

Here we used movie-watching fMRI data from 176 HCP subjects to map a novel cortical brain network involved in the face processing. This network was comprised of four distinct face patches (FFPs) located in the dorsolateral prefrontal cortex. FFPs were functionally connected to mSTS and responded to facial and social cues in the naturalistic movies. We showed that elevated activation of FFPs was linked to improved performance in a social cognition task. Furthermore, we identified two groups of subjects based on the topological organization of face-selective patches and assessed the structural and functional differences across these two subject groups.

Functional localizer tasks with static stimuli do not consistently identify all face-processing regions. Specifically, high-level face-selective regions may be less active in these tasks due to the lack of contextual, social, and semantic information in the stimuli [9, 30, 74]. A naturalistic audiovisual movie-watching task is a more powerful paradigm for studying the face-processing network, offering a closer approximation of real-world scenarios [40]. The movie stimulus is rich in sensory aspects, is cognitively engaging, and contains dynamic faces, facial emotions, and human-specific social interactions. Using a movie-watching task and a multivariate connectivity analysis, we revealed a more detailed organization of face-selective cortical network encompassing frontal regions. This network is activated during movie watching to process faces and integrate facial information with multiple other sources of information (e.g., social, linguistic, semantic, and emotional content) [12, 17, 58, 97, 98, 101].

The social processing network in the brain encompasses regions including temporo-parietal junction (TPJ), superior temporal sulcus (STS), and medial prefrontal cortex [19, 51, 69, 83, 99]. Although the role of the lateral prefrontal cortex in social processing is less explored in humans, studies in non-human primates have shown activation in this region during social perception tasks [84, 89, 90], as well as its connectivity with the core face processing system [87]. A recent work by Rajimehr et al. [72], using a movie-watching paradigm, identified a social cognition network which included regions in the lateral prefrontal cortex in the right hemisphere. In this study, we showed that within this frontal social network, patches linked to face processing (FFPs) are present. This network may also contain body-selective patches, considering the fact that social perception involves understanding both facial and bodily physical interactions. The presence of such patches in the lateral prefrontal cortex could be tested in future studies.

Our multivariate connectivity maps showed strong connectivity between FFPs and the right mSTS face patch located within TPOJ. Previous studies have shown that mSTS is involved in social face processing [51], and is activated during the interpretation of facial movements as opposed to object motion [107]. This region also shows marked preference for audio speech over other types of non-speech vocal sounds [20]. Thus, the connectivity with mSTS implies the involvement of FFPs in social processing. Furthermore, using a reverse-correlation analysis, we found that FFPs were highly responsive to movie frames featuring human presence (“man” and “woman”) and communicative actions (“talk”), suggesting their role in processing social face cues. Based on previous studies in monkeys, face patches in the frontal cortex could also be implicated in the processing of facial expressions [77, 95]. However, a direct comparison between FFPs and the emotional face activations showed that the hotspots of activity in the emotion task were mostly outside of FFPs (see Fig. S5).

The examination of individual-level cortical connectivity maps of the mSTS showed two distinct subject groups based on the spatial organization of FFPs. The most dorsal and most ventral FFPs (FFP1 and FFP4, respectively) were present in the same location in both subject groups, whereas FFP3 and FFP2 were present only in G1 subjects (*N* = 132) and G2 subjects (*N* = 44), respectively. This intriguing distinction led us to ask whether the two groups are more fundamentally distinct in their cortical organization. Upon examining each group’s cortical folding map, we observed anatomical differences between the groups in the vicinity of FFP2 and FFP3. This finding concurs with the previous research, indicating that the location of functional activity in the brain is dictated by underlying sulcal pattern [57, 63]. Furthermore, when assessing activations in the social cognition task, we found that G1 subjects showed greater activation in FFP3, while G2 subjects showed greater activation in FFP2. This activation disparity, assessed from the same cohort but through an fMRI task of social cognition, provides further evidence for the possible role of FFPs in social processing.

The stratification of subjects into two subtypes in this study challenges the conventional reliance on group-average “norm” maps. While group-average maps are informative, they can wash out individual differences and unique individuals’ characteristics. However, studying brain maps at the level of individual subjects can bring uncertainty about discerning noise from genuine biological variations across subjects [23]. Subtyping subjects can be an intermediate step that fills the gap between generalized and idiosyncratic organization of brain maps [22, 54] and can reveal layers of information otherwise obscured. Specifically, subtyping can be useful in functional and structural studies of the frontal cortex where considerable individual differences exist [43, 53]. In this regard, our study revealed a clear distinction in the functional and structural organization of FFPs between two subject groups. These two groups may also have distinct genetic profiles, which could be examined in future studies.

A subsequent question is what makes each of the four FFPs distinct in terms of functional properties. We provided evidence for specific contributions of FFPs to social face processing based on the partial correlation analysis. FFP1 was selectively connected to areas involved in visual motion processing (MST), eye movement control (FEF and SCEF), and social processing (55b, PSL), suggesting a role for FFP1 in processing dynamic social-facial cues to guide eye movements. FFP2 was selectively connected to mid-level visual and scene-selective areas, hence, we hypothesized that FFP2 was involved in integrating facial information with the background scene context. FFP3 was selectively connected to auditory cortex, hence we hypothesized that FFP3 was involved in integrating facial and linguistic information. FFP4 was selectively connected to regions in the default mode network, suggesting a role for FFP4 in social-semantic processing of faces. FFP4 was also connected to areas 44 and 45 in the right hemisphere, which are part of the social processing network in the brain [71, 72]. These hypotheses regarding the function of the FFPs are derived based on their unique connectivity patterns and should be rigorously re-evaluated in future controlled studies.

We provided further evidence for the role of FFPs in social cognition by testing the relationship between the subjects’ performance in the HCP social cognition task and the level of activations in FFPs in a larger cohort of subjects (*N* = 1, 048). Our analysis showed a strong correlation between activations in FFP1, FFP3, FFP4 and behavioral performance in the social cognition task, backing up the role of FFPs in social cognition. We did not find a significant behavioral difference between the two subject groups. However, given the inter-subject variations in the spatial organization of FFPs, there might be behavioral consequences when watching movies and understanding their narratives. For instance, subjects having FFP2 may be more focused on perceiving the visual content of movies, while subjects having FFP3 may be more focused on tracking the conversational content of movies. Such differences in movie-watching strategies, if verified, could have important implications for the film industry and production.

The present findings should be considered with respect to some methodological limitations. First, the eight classic face-selective areas were identified using a conservative thresholding approach, selecting 1% of vertices with the highest response for the contrast of faces vs. other categories. This approach identified mSTS only in the right hemisphere. Since the multivariate connectivity maps were constructed for classic face-selective areas and the ipsilateral hemisphere, FFPs were identified and assessed exclusively in the right hemisphere. FFPs could also be present in the left hemisphere, though based on the previous literature, it is expected to have a right-hemisphere dominance for the face processing [18, 26]. Second, the inference about the role of individual FFPs is based on correlational findings and requires further validation through causal studies. Third, as seen in the multivariate connectivity maps of MFA and pSTS (Fig. S2), face-responsive regions may also exist in the medial prefrontal cortex. These regions should be explored in future studies.

## METHODS

### Subjects

For working memory, social cognition, emotion processing, and language processing tasks, we used the S1200 data release from the Human Connectome Project (HCP). For the movie-watching task, we used the “HCP-7T” dataset which is a part of the HCP-S1200 data release. The movie-watching dataset included data from 184 subjects; however, only 176 subjects (106 females, 70 males) had complete functional data for movie-watching scans and hence were included in the analysis.

Subjects in all tasks were healthy young adults aged 22–35 years old, and were scanned at Washington University in St. Louis (3T data) and the Center for Magnetic Resonance Research at the University of Minnesota (7T data). All subjects had normal or corrected-to-normal visual acuity. The HCP data were acquired using protocols approved by the Washington University institutional review board, and written informed consent was obtained from all subjects [104].

### Structural MRI data acquisition

The structural MRI data was acquired using a customized 3 Tesla Siemens Skyra scanner with a standard Siemens 32-channel RF-receive head coil. For each subject, at least one 3D T1-weighted MPRAGE image and one 3D T2-weighted SPACE image were collected. The parameters of the T1-weighted MPRAGE sequence were as follows: TR/TE/TI = 2400*/*2.14*/*1000 ms, flip angle = 8^◦^, bandwidth = 210 Hz*/*pixel, field of view = 224 *×* 224 mm, spatial resolution = 0.7 mm^3^, GRAPPA factor = 2, echo spacing = 7.6 ms. The acquisition time was 7 minutes and 40seconds. The parameters of the T2-weighted SPACE sequence were as follows: TR/TE = 3200*/*565 ms, flip angle = variable, bandwidth = 744 Hz*/*pixel, field of view = 224 *×* 224 mm, spatial resolution = 0.7 mm^3^, GRAPPA factor = 2. The acquisition time was 8 minutes and 24 seconds.

### Functional MRI data acquisition

#### Working memory, social cognition, language processing, and emotion processing tasks

The task fMRI data was acquired using a customized 3 Tesla Siemens Skyra scanner with a standard Siemens 32-channel RF-receive head coil. Data was collected using a slice-accelerated, multi-band gradient-echo echo planar imaging (EPI) sequence with the following parameters: TR/TE = 720*/*33.1 ms, flip angle = 52^◦^, bandwidth = 2290 Hz*/*pixel, field of view = 208 *×* 180 mm, spatial resolution = 2.0 mm^3^, number of slices = 72, multiband factor = 8, and echo spacing = 0.58 ms. The direction of the phase encoding altered between left-to-right and right-to-left across runs.

#### Movie-watching task

The movie-watching fMRI data was acquired using a 7 Tesla Siemens Magnetom scanner with a Nova 32-channel RF-receive head coil. Data was collected using an EPI sequence with the following parameters: TR/TE = 1000*/*22.2 ms, flip angle = 45^◦^, bandwidth = 1924 Hz*/*pixel, field of view = 208 *×* 208 mm, spatial resolution = 1.6 mm^3^, number of slices = 85, multi-band factor = 5, and echo spacing = 0.64 ms. The direction of phase encoding altered between posterior-to-anterior and anterior-to-posterior across runs.

### Experimental paradigms

#### Working memory task

The working memory task was a version of the *N*-back task to assess working memory and cognitive control [8]. By presenting blocks of trials that consisted of pictures of faces, places, tools, and body parts, this task could also be used as a “functional localizer” to obtain category-specific representations. Subjects performed two runs of the working memory task. Each run contained eight task blocks (25 s each) and four fixation blocks (15 s each). The four different stimulus types (faces, places, tools, and body parts) were presented in separate task blocks. Each task block contained ten trials. On each trial, the stimulus was presented for 2 s, followed by a 500 ms inter-trial interval. Within each run, four blocks used a 2-back working memory task (respond “target” whenever the current stimulus was the same as the one 2-back) and the other four blocks used a 0-back working memory task (respond “target” whenever the current stimulus was the same as the target stimulus presented at the start of the block). A 2.5 s cue indicated the task type (and target for 0-back) at the start of the block. In each block, there were two targets and 2–3 non-target stimuli (repeated items in the wrong *N*-back position, either 1-back or 3-back).

#### Social cognition task

The social cognition (theory of mind) task consisted of brief 20-second video clips sourced from Castelli et al. [13] and Wheatley et al. [100], featuring geometric shapes (squares, circles, triangles) with either deliberate interactions or random motion patterns on the screen. Clips were separated by 15-second fixation blocks. After each clip, subjects were asked to discern whether the shapes’ movements implied a social/mental interaction or had no observable interaction (random movements). If subjects were not sure about the existence of interaction between the shapes’ movements, they could indicate uncertainty by choosing a “Not Sure” option. Each subject participated in two runs encompassing five video blocks—in one run, two video blocks included mental interaction and three included random motion; in the other run, three video blocks included mental interaction and two included random motion.

#### Language processing task

The language task consisted of auditory blocks of story and math with varying lengths (average of approximately 30 seconds), originally developed by Binder et al. [10]. During story blocks, subjects listened to auditory narratives (5–9 sentences) from Aesop’s fables, followed by a 2-choice question regarding the story’s topic. During math blocks, subjects were auditorially presented with a series of simple arithmetic operations (addition and subtraction), followed by 2-choice questions. The math task was adaptive to try to maintain a similar level of difficulty across subjects. Each subject participated in two runs encompassing interleaved four blocks of story task and four blocks of math task.

#### Emotion processing task

The emotion processing task was adapted from the one developed by Hariri et al. [38]. This task consisted of blocks of trials including images of faces with angry or fearful expression, or shapes as the control for facial emotion processing. At each task trial, subjects were asked to decide which of the two stimuli in the bottom of the screen match the one on top using a finger button press. Each subject participated in two runs encompassing six 21-second blocks (3 face blocks and 3 shape blocks), with 8 seconds of fixation at the end of each run. Each block was preceded by a 3-second task cue (“shape” or “face”). Within each block, six 2-second trials of the same stimulus type (shape or face) were presented with an inter-trial interval of 1 second.

#### Movie-watching task

The movie-watching task consisted of four functional runs where subjects passively viewed a series of audiovisual movie clips, each lasting approximately 15 minutes. Each run consisted of 4 or 5 clips which varied in duration from 1:03 to 4:19 (minute:second). Before the first movie clip, in between clips, and following the last clip, a 20-second period of rest existed, which was indicated by the word “REST” displayed in white text on a black background. The first and third runs contained clips from independent films (both fiction and documentary), which were made available under a Creative Commons license on Vimeo. The second and fourth runs contained clips from Hollywood films prepared by Cutting et al. [16]. The last clip of all runs was identical and included a montage of brief (1.5 s) videos. A brief description of each clip can be found in Finn and Bandettini [28]. The audio was delivered via Sensimetric earbuds, and movies were presented in full-screen mode, occupying a visual angle of 21.8^◦^ in width and 15.7^◦^ in height.

### Analysis of structural MRI data

Structural images (T1-weighted and T2-weighted) were processed using the Human Connectome Project (HCP) three-stage processing pipeline. The first step (“PreFreeSurfer” pipeline) corrected for geometric distortions and bias fields in structural images, aligned T1-weighted and T2-weighted images, created the initial brain mask, and registered the subject-specific space to MNI152 standard space using nonlinear volume-based registration. The second step (“FreeSurfer” pipeline; based on modified FreeSurfer version 5.2) reconstructed the white and gray matter surfaces and the subcortical mask using high-resolution (0.7 mm) T1-weighted and T2-weighted images and applied folding-based registration to align individual surfaces with the standardized fsaverage template. The third step (“PostFreeSurfer” pipeline) mapped cortical data to Conte69 surface template and converted FreeSurfer outputs to standard NIFTI and GIFTI file formats. Registered cortical vertices were combined with subcortical gray matter voxels to construct the standard “CIFTI grayordinates” space, comprising 91, 282 vertices/voxels with approximately 2 mm cortical vertex spacing and 2 mm isotropic voxels [33].

### Curvature maps

Curvature maps highlight the geometry of cortical folding patterns, with negative values representing sulci (inward folds) and positive values representing gyri (outward folds). For each point (vertex) on the cortical surface at the gray matter-white matter boundary, cortical curvature was quantified using mean curvature measurement—the average of principal curvatures derived from the inverse radius of the osculating circles at that point. The curvature maps were taken directly from FreeSurfer and subsequently mapped onto the standard CIFTI grayordinates surface space [35].

### Analysis of functional MRI data

Functional data were preprocessed using the HCP minimal preprocessing pipelines described by Glasser et al. [33]. Preprocessing steps included the removal of spatial artifacts caused by the B0 field inhomogeneity and gradient nonlinearity, field map-based unwarping of EPI images, motion correction, brain-boundary-based registration of EPI data to structural MRI scans, non-linear registration to MNI152 standard space, and grand-mean intensity normalization. Next, the gray matter data were projected onto the standard grayordinates space, and minimally smoothed using a 2 mm full width at half maximum (FWHM) Gaussian kernel. Notably, mapping fMRI data to the standard grayordinates space was performed using multi-modal areal-feature-based surface registration (MSMAll) which enhances alignment across subjects by incorporating folding patterns, myelin maps, resting-state networks, and visuotopic connectivity maps during registration [34, 35, 75].

For the task-based fMRI analysis, the preprocessed functional time-series were entered into a modified FSL’s FEAT analysis pipeline. FSL’s FILM algorithm was used to compute the first-level task fMRI statistics, and FSL’s FLAME algorithm was used in a fixed-effects analysis to combine data across runs within subjects [8]. In this study, we employed the beta maps of contrasts provided by the HCP.

For the movie-watching fMRI analysis, data were cleaned up for artifacts and structured noise using sICA+FIX. Minimal high-pass filtering with a cutoff of 2000 s was also applied. The effect of this filter was similar to linear detrending of the fMRI signal.

### Classic face-selective regions

To localize the face-selective regions, we used the category localizer data from the *N*-back working memory task. We disregarded the effect of memory load by collapsing the fMRI data for each category (face, body, place, and tool) across 0-back and 2-back conditions. For the contrast of interest (faces vs. all other categories), fixed-effects analyses were conducted to estimate the average effects across runs within each subject, then mixed-effects analyses were conducted to obtain the group-average map. To identify the classic face-selective regions, we thresholded the group-averaged beta coefficient map to select one percent of vertices/voxels in the CIFTI grayordinates space (913 out of 91, 282 vertices/voxels) with the largest response in the contrast of faces vs. other categories. This thresholding approach resulted in the identification of eight cortical face-selective regions [1]. Among the eight face-selective regions, five were in the right hemisphere (FFA, OFA, MFA, pSTS, and mSTS), and three were in the left hemisphere (FFA, MFA, and pSTS) (Figure 1C,D).

### Multivariate connectivity analysis

We implemented a multivariate analysis approach combining Partial Least Squares Regression (PLSR) with surface-based searchlight analysis to identify ipsilateral cortical regions functionally connected to core face-processing areas during movie-watching [2, 3, 60]. For each searchlight ROI (including 100 vertices), a PLS model was trained to predict the mean time-course of activity in a face-selective target area (*y*; 1*×* number of time-points) based on the activity pattern across ROI vertices (*X*; 100*×* number of time-points). The PLS latent variables captured the shared covariance between ROI’s time-courses (*X*) and target area’s mean time-course (*y*). For each model, we assessed predictive accuracy on held-out test data using “explained variance” (Eq. 1), which quantifies the similarity between actual (*y*) and predicted (*ŷ*) time-courses. This value was assigned to each searchlight ROI’s central vertex, and represented that vertex’s multivariate connectivity with the face-selective target area (see Supplementary Figure S1) [105].

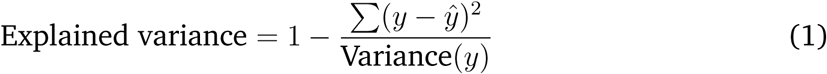

For the PLS analysis, we used movie-watching fMRI data across all four runs. Each run included several short movie clips (run1: 5 clips, run2: 4 clips, run3: 5 clips, run4: 4 clips). We discarded data from rest periods between clips as well as the initial 4 seconds of each clip.

Optimizing the PLS model required determining the appropriate number of latent components (ranging from 1 to 100) for each searchlight ROI. Therefore, for each ROI, 100 PLS models with varying component numbers should be constructed using train data, and models’ performance should be assessed on test data to specify optimal parameter value. This optimization process would ideally be performed for each of the 176 subjects across approximately 30, 000 searchlight ROIs (within the ipsilateral hemisphere) when predicting the mean time-course of each of the eight face-selective areas. To manage this substantial computational load, the analysis was conducted in two stages.

In the first stage, eight representative subjects were selected from the full cohort of 176 individuals to determine an optimal component number (from the range of 1 to 100) as the parameter for PLS models. For each of the four movie runs, we implemented a leave-one-clip-out cross-validation procedure, where each clip sequentially served as test data while the remaining clips from that run constituted the training data. For each fold, we constructed PLS models with varying numbers of components (1–100) and identified the component number that maximized prediction accuracy on the test clip. This process generated 18 optimal component values (corresponding to the total number of clips across all runs) for each subject, target area, and searchlight ROI combination. We then computed the median of these values across all eight subjects, eight target areas, and approximately 30, 000 searchlight ROIs, yielding an optimal value of 8 components.

In the second stage, we applied the chosen parameter of 8 components to construct PLS models for all 176 subjects. The leave-one-clip-out cross-validation procedure was used in this stage as well. The maps from all 176 subjects were subsequently used to obtain the group-average multivariate connectivity map (Cohen’s d) for each target area. The multivariate connectivity maps for the eight face-selective areas are shown in Supplementary Figure S2.

### Localization of FFPs

The right mSTS multivariate connectivity map revealed a distinct connectivity pattern within the lateral prefrontal cortex (Figure 1E). We created a mask of prefrontal regions with enhanced mSTS connectivity by thresholding the group-average Cohen’s d map at 0.7, resulting in an area of 2,032 vertices (Figure S2A). Within this mask, we consistently observed three patches with strong connectivity to the right mSTS across subjects. While the dorsal and ventral patches showed stable localization across individuals, the central patch showed variability in its location (Supplementary Figure S3). To characterize this spatial variability across subjects, we used the Isomap dimensionality reduction technique with the parameter *k* set to 8. In this approach, each vertex within the mask was treated as an individual feature, resulting in 2, 032 features per subject. Isomap reduced this high-dimensional feature space to a two-dimensional representation (Figure 2C).

We then used DBSCAN (Density-Based Spatial Clustering of Applications with Noise) to classify subjects based on the spatial organization of FFPs within the prefrontal mask. DBSCAN method of clustering relies on two parameters: epsilon (eps) and minimum points (minPts). We systematically explored parameter combinations across a range of values, with eps varying from 2 to 5 in increments of 0.1 and minPts varying from 10 to 20 in increments of 1. We focused on clustering solutions that produced exactly two clusters and applied majority voting to assign a definitive cluster label to each subject. This methodology revealed two subject groups characterized by distinct mSTS-prefrontal connectivity profiles: a larger group (*N* = 132) exhibiting a strong connectivity between mSTS and FFP3 and a smaller group (*N* = 44) exhibiting a strong connectivity between mSTS and FFP2 (Figure 2D).

To further assess the structural and functional differences between the two subject groups, we needed to define a boundary for each FFP. We used a local thresholding approach to delineate these boundaries. First, we identified the vertices with maximum connectivity value for each of the six visible patches (FFP1, FFP3, and FFP4 in G1, and FFP1, FFP2, and FFP4 in G2). We then set a threshold at 75% of the mean of these six maximum values. The patches were defined by selecting vertices which had connectivity values greater than the threshold value. This approach did not impose geometric constraints on the patches (e.g., a circular shape).

## Data analysis software

The HCP data underwent preprocessing using the publicly available HCP pipelines. Our analyses utilized various software packages, including Connectome Workbench command-line tools (http://www.humanconnectome.org/software/connectome-workbench.html), FreeSurfer [29], FSL [47], and Matlab. The Permutation Analysis of Linear Models (PALM) software [102] was used for the TFCE analysis. We employed the “wb_view” GUI in Connectome Workbench to visualize the cortical brain maps.

## Data/code availability

The movie-watching and task fMRI data are part of the publicly available and anonymized HCP database (https://www.humanconnectome.org). Multi-modal cortical parcellation files are available for download in the BALSA database (https://balsa.wustl.edu/study/show/RVVG). All analysis codes are available for sharing upon request.

## Acknowledgments

We thank members of the HCP team for helpful advice during data analysis. This research was supported by Institute for Research in Fundamental Sciences (IPM).

---

**Figure S1.**
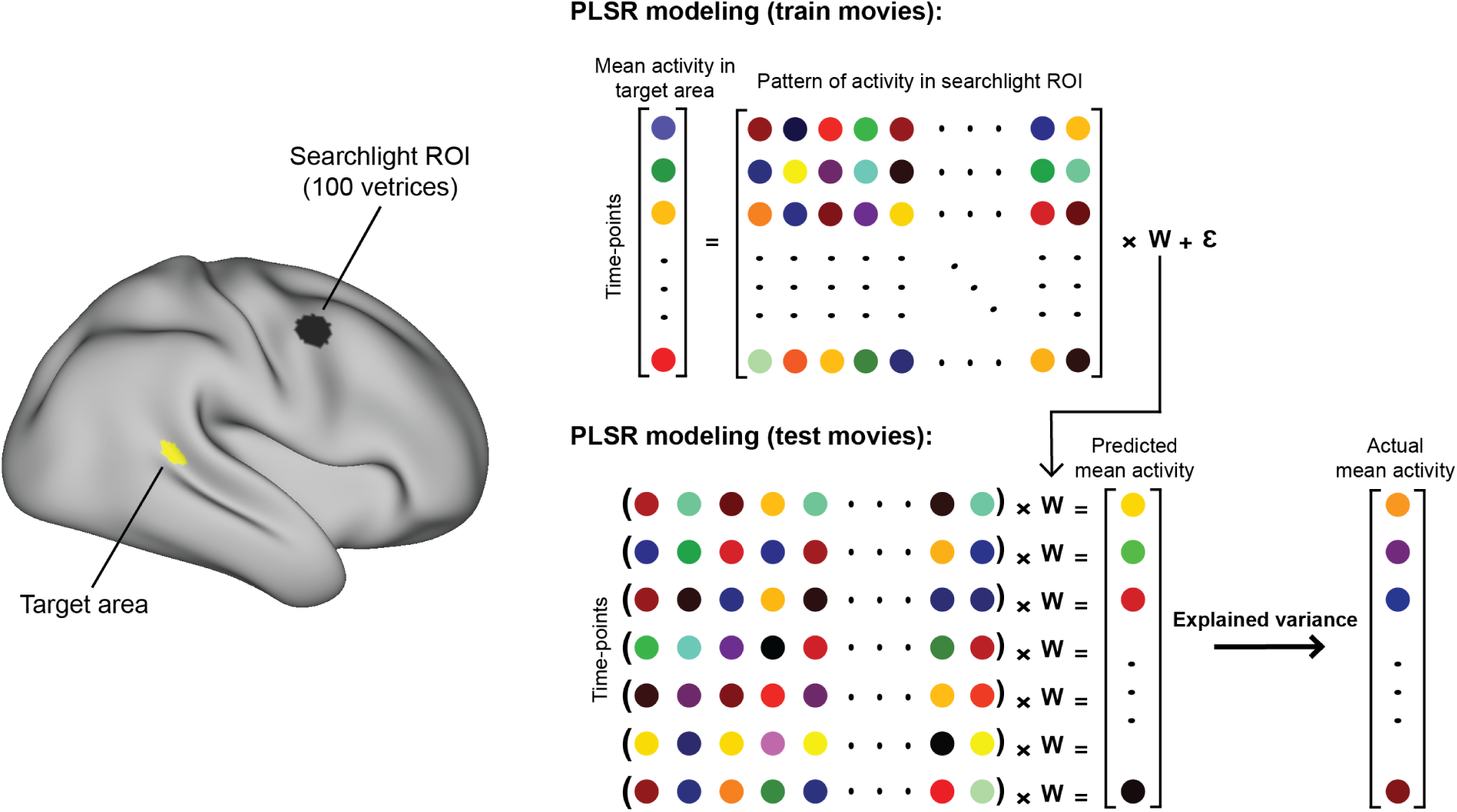
Multivariate connectivity analysis. We used surface-based searchlight ROIs to identify cortical regions where activity patterns explain the mean time-course of activity in a target brain area. For each searchlight ROI, a PLS model is trained using a subset of data, then the trained model is applied to the test data to predict the mean time-course of activity in the target area. The similarity between predicted and real time-courses is quantified by explained variance. The explained variance value, indicating multivariate connectivity between the target area and the searchlight ROI, is assigned to the central vertex of the ROI.

**Figure S2.**
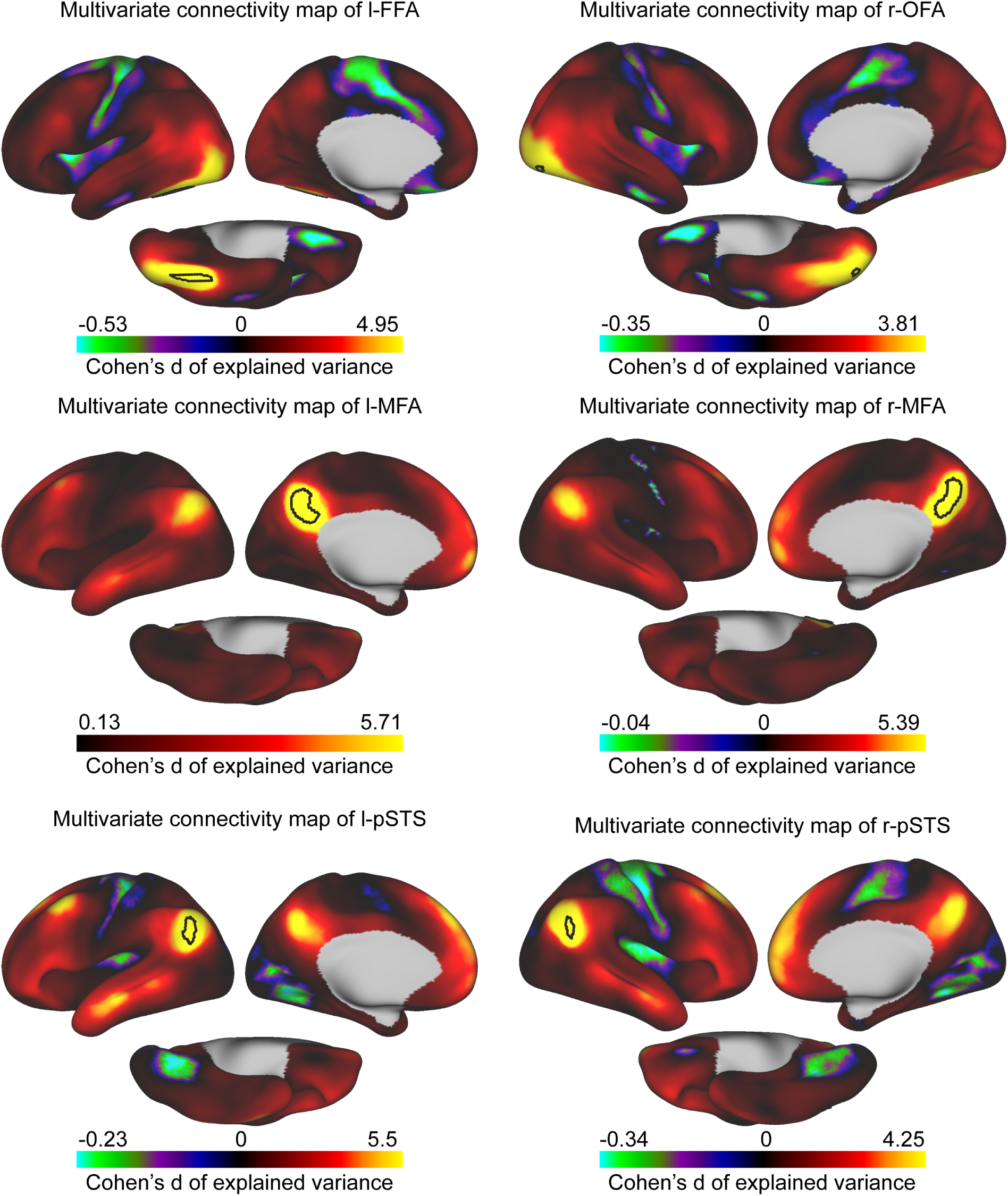
Group-average multivariate connectivity maps of classic face-selective regions (black outlines) with ipsilateral cortical regions. Brain cortical maps are shown on lateral, medial, and ventral views of inflated fs-LR surface.

**Figure S3.**
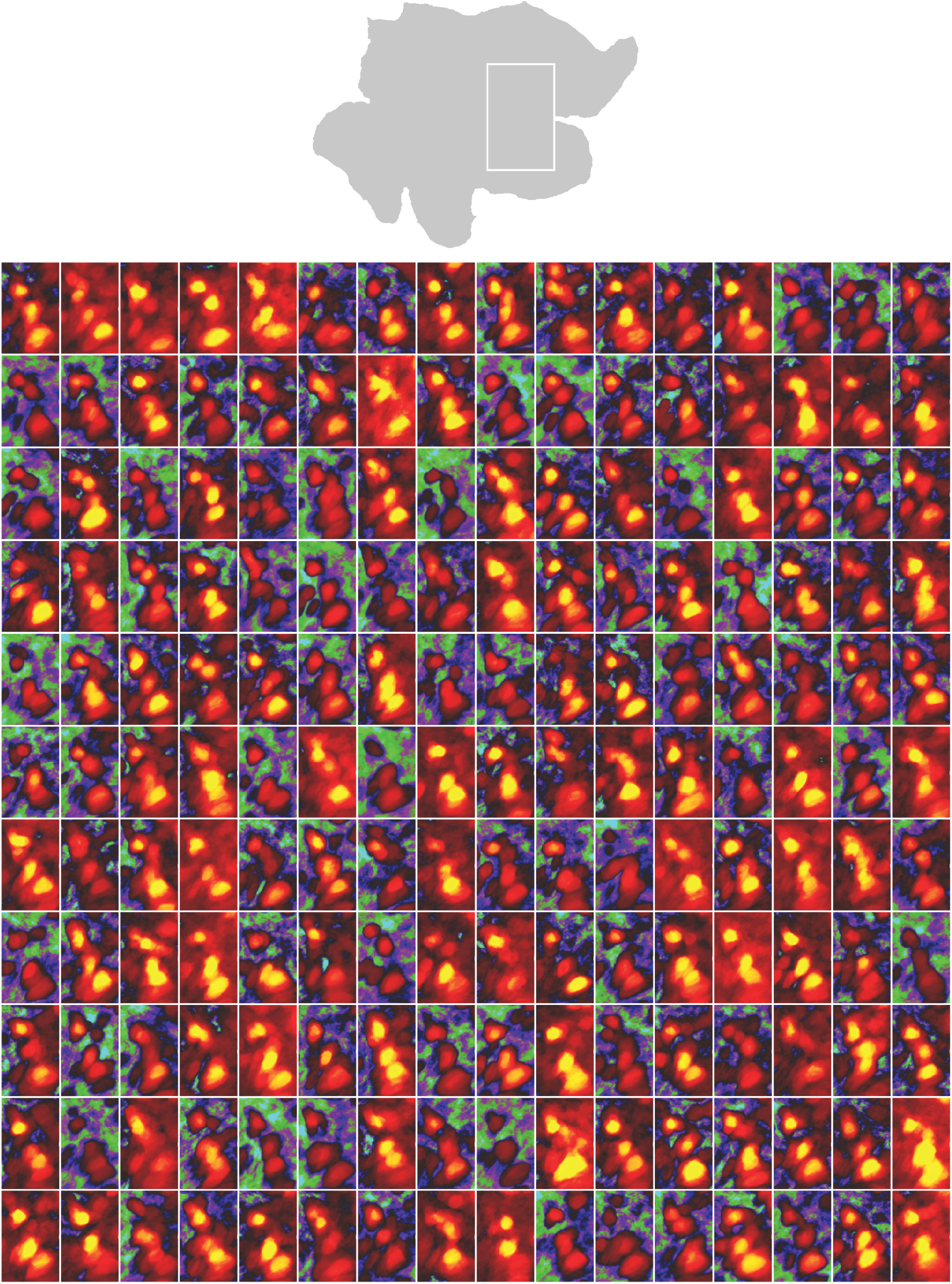
Organization of FFPs in individual subjects. For each subject, the multivariate functional connectivity of right mSTS is obtained. The white outline on the right hemisphere’s flattened fs-LR surface highlights a square region magnified in the individual subjects (*N* = 176) to show the variation in the topology of patches in the frontal cortex.

**Figure S4.**
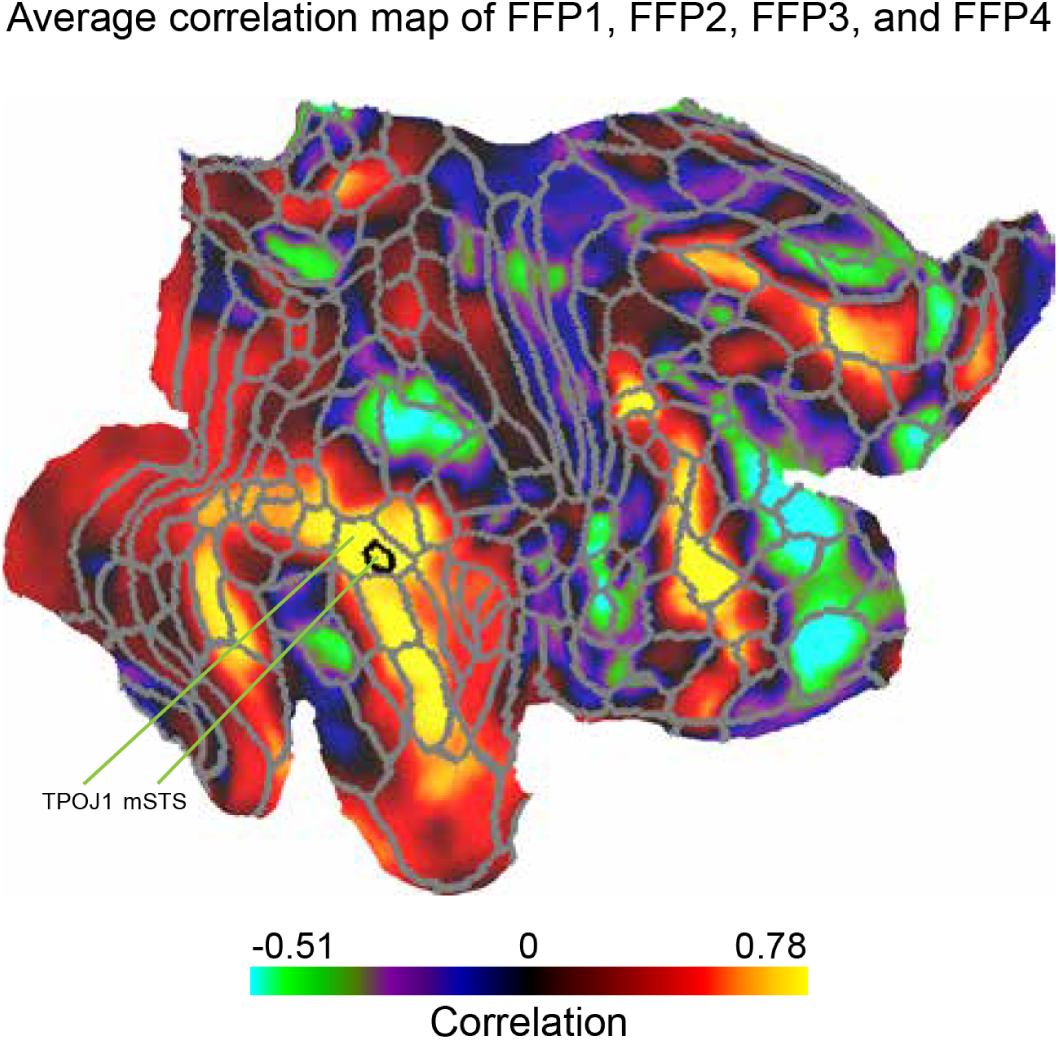
Univariate connectivity between the mean time-course of activity in all FFPs and the time-courses of all cortical vertices in the right hemisphere. Brain cortical map is shown on the right hemisphere’s flattened fs-LR surface. Borders of multi-modal cortical parcellation [34] (in gray) and the mSTS face-selective area (in black) are displayed on the surface map. The temporo-parietal junction (TPOJ1 parcel) is the region with the highest univariate connectivity to FFPs.

**Figure S5.**
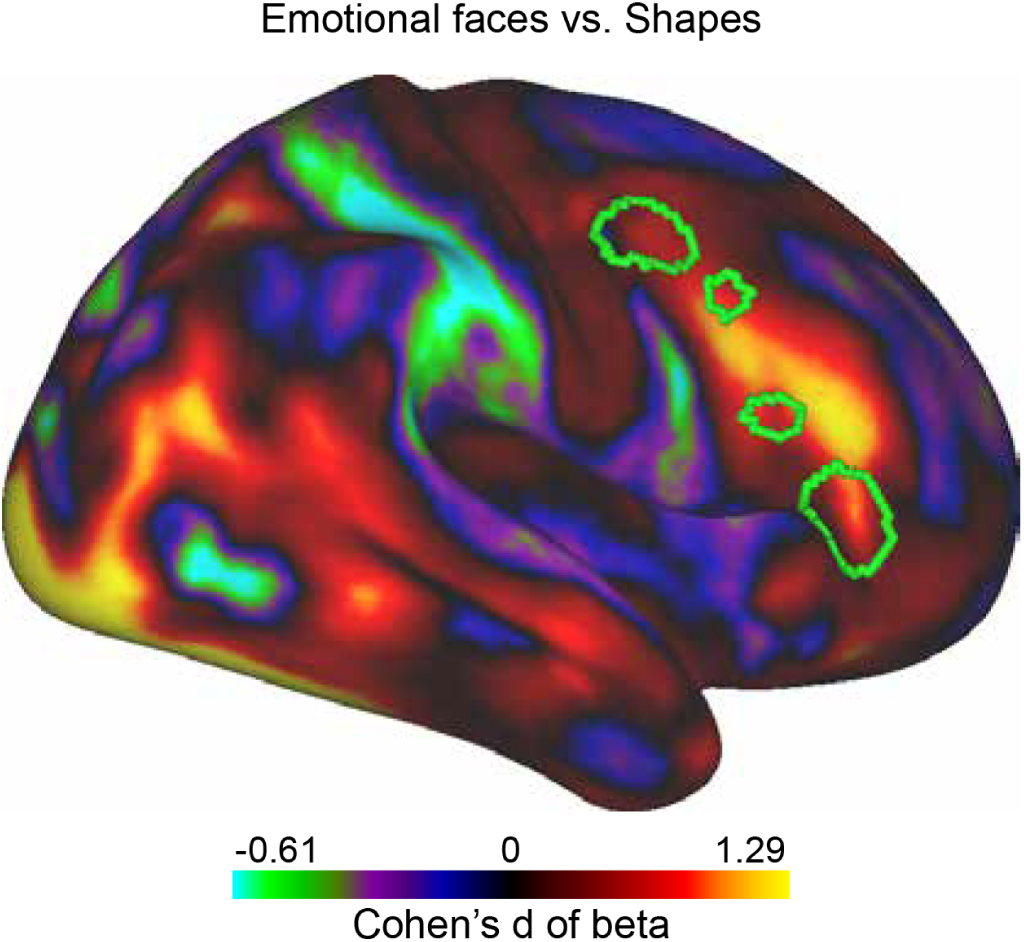
Group-average Cohen’s d effect size map for the contrast of emotional faces versus shapes. The map is obtained from the HCP emotion processing task. Green borders indicate FFPs.

